# E-PoSa: a novel and effective tool for sampling pollen directly from flowers

**DOI:** 10.1101/2023.06.30.547178

**Authors:** E. Pioltelli, L. Guzzetti, L. Tonietti, A. Copetta, P. Biella, L. Campone, A. Galimberti

## Abstract

1. Pollinator insects are declining worldwide also due to the alteration of their diet with severe implications on their health status. Pollinators diet relies mainly on flower rewards (i.e., pollen and nectar) and a precise characterization of their chemical composition is crucial in defining pollinators’ nutritional ecology. In this context, the pollen represents a challenging source to investigate, especially due to operative challenges during collection operations and to the small amounts produced per flower.
2. Here, we designed and tested a novel, easy-to-assemble tool for pollen sampling: E-PoSa (Electronic Pollen Sampler), based on the use of a portable vacuum cleaner. We compared it with some of the most used sampling methods for pollen (i.e., anthers sieving and sampling of the whole anthers) by looking at the differences in their quantitative recovery and nutritional profile. Its applicability in ecological studies was also corroborated by an assessment of its recovery rate obtained from a panel of wildflowers species in an operational environment.
3. The data obtained showed a significantly higher pollen recovery capacity of E-PoSa compared to the conventional sieving approach and the success in retrieving enough pollen to conduct phytochemical analyses from a broad range of flower morphologies in the field. Our results also demonstrated that high purity pollen can be collected with E-PoSa and that the device does not introduce any significant variation in the nutritional analysis compared to the conventional sieving.
4. This new sampling approach represents a cheap and easy-to-assemble tool encouraging its future use not only in the field of pollen nutrition but also in a wide variety of other contexts related to pollination ecology. Acknowledging the potential influences of the sampling techniques and moving towards shared standardized field protocols will advance the comprehension of species interactions and foraging patterns of pollinators and their nutritional needs.

## 1. Introduction

Insect pollinators are declining worldwide due to multiple global issues such as climate change (Vasiliev D. & Greenwood S., 2021), exposure to pesticides (Goulson et al., 2015), and habitat loss (Potts et al., 2016). The depletion of dietary resources (i.e., pollen and nectar) due to the loss of flower-rich habitats and/or their contamination due to the use of agrochemicals, is one of the main risk factors for pollinators (Vaudo et al., 2015; Hülsmann et al., 2015). Considerable evidence about the importance of adequate nutrition for pollinators conservation have fostered a growing interest in the investigation of the nutritional landscape for a better understanding of the relationships existing between pollinating insects and floral resources (Leonhardt et al., 2022; Vaudo et al., 2018; Vaudo et al., 2016; Venjakob et al., 2022). Many pollinators feed on pollen which represents the main protein source (Nicolson et al., 2018) and in this context, nutritional analyses of pollen are of paramount importance. However, these studies are often challenging at the analytical level, due for example to the scarcity of collected material or its frequent contamination by other floral parts (e.g., anthers and petal parts). Indeed, many flowers produce a low amount of pollen (< 1 mg; Jeannerod et al., 2022).

Researchers have developed many sampling approaches to improve sampling efficiency and to face the critical issue of retrieving an adequate quantity of pollen to achieve reliable nutritional analyses (Jeannerod et al., 2022; Kendel and Zimmermann 2020; Knäbe et al., 2015). This wide panel of pollen sampling results in a great heterogeneity among the different studies, hampering the comparison between the results obtained. With such a variety of available methodologies for the sampling of floral resources, each one with different benefits and drawbacks, the need to standardize the collection effort is becoming even more urgent.

Here, we propose a novel non-invasive tool for pollen sampling. The device, E-PoSa, (Electronic Pollen Sampler) is based on the use of a commercially available portable vacuum specifically adapted to this purpose and allows the collection of highly pure pollen grains directly from the flower in a non-destructive way. To validate this new approach, we provide a comparison with some of the most common methodologies adopted for pollen sampling to evaluate the magnitude of the differences that occur in the recovery efficiency and in the subsequent analysis of the nutrient composition.

## 2. Material and Methods

### 2.1. Assemblage and use of E-PoSa

To conduct a study on the pollen nutritional composition of wildflowers in northern Italy, we developed an effective and easy-to-assemble device that allows the collection of pollen with a yield suitable for chemical characterization. E-PoSa is based on the use of a portable vacuum cleaner with a removable plastic mask. The tool is made up of the following parts: a portable vacuum, a 5 ml tube with the head cut and the lid drilled; an inox mesh sheet, a paper filter, and laboratory film (Figure 1). The first step for assembling the device is preparing the 5 mL tube. The head of the tube needs to be cut approximately 0.5 - 1 cm from the tip using a sharp knife (Video S1). Second, the tube lid must be separated and drilled by using a conical drill bit (Video S1). The next step is cutting the paper filter with a diameter slightly larger than that of the test tube cap. Then, close the tube by paying attention that the paper filter remains in the correct position by completely covering the hole previously made on the cap (Video S1). After cutting a small square from the inox mesh sheet, heat it quickly using a lighter, place it at the top of the tube, and hold it in place for a few seconds to ensure it does not detach (Video S1). The last step is to connect the tube to the head of the vacuum and secure it by using the laboratory film (Video S1). The time required for the assembly is estimated to be less than 10 minutes. A few seconds are required for re-placing the pre-assembled collection tubes. The battery life of the portable vacuum used in this study is about 30 minutes, but it can be easily extended by integrating it with a portable power bank that allows its use also in remote sites. Details regarding the materials utilized and their costs are reported in Table S1.

**Figure 1:**
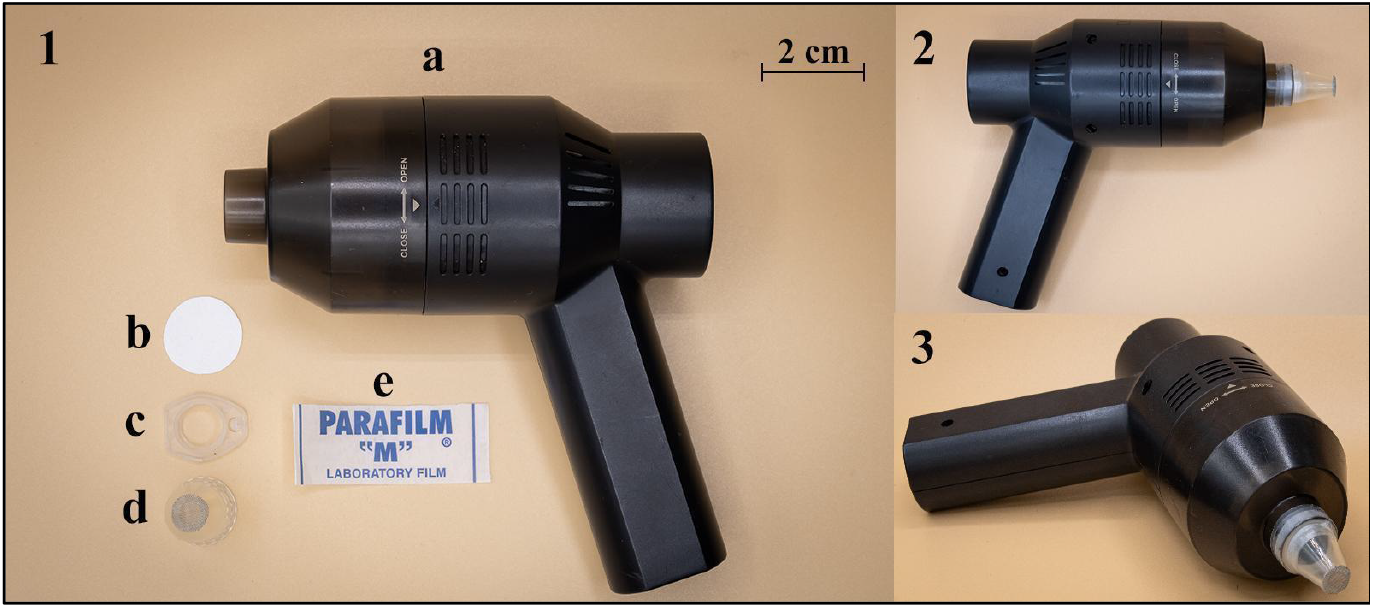
Assemblage of the E-PoSa. Overview of all the materials needed for the assembly of the E-PoSa device (a. Portable vacuum; b. Paper filter with 25 μm pore size; c. 5 mL Eppendorf tube’s cap drilled; d. 5 ml Eppendorf tube with Stainless steel mesh with 75 μm mesh size at the tip; e. strip of parafilm); 2. Top view of the fully assembled E-PoSa 3. Frontal view of the fully assembled E-PoSA.

The use of E-PoSa does not require expertise. The pollen can be collected by simply turning on the vacuum and moving it on the flowers to be sampled. Pollen is aspirated and accumulates on the paper filter. Due to the transparency of the tube, the amount of pollen collected can be easily estimated by eye and once enough pollen is collected, it can be directly transferred to another tube by removing the inox mesh from the tip of the E-PoSa and gently tapping on the bottom of the adapted tube. The system is useful with flowers of different morphology. In the case of small flowers or flowers that have anthers within the corolla (e.g., Lamiaceae or Fabaceae) a 20 μL tip can be attached to the tube and fixed with parafilm to allow a more accurate and effective pollen sampling.

### 2.2. Study species

The target flower species were selected on their taxonomy to account for a wider set of families characterized by different floral morphologies and different amounts of pollen produced. For the collection of anthers and pollen grains, a panel of three species was selected: *Tropaeolum majus* L. (Fam.: Tropaeolaceae), *Hippeastrum vittatum* Herb. (Fam.: Amaryllidaceae), *Alstroemeria aurea* Graham (Fam.: Alstroemeriaceae) (Fig. 2 A, B, C). The flowers were covered with a nylon mesh 24 h before sampling to avoid possible depletion of resources by pollinator visits. The study took place at the C.R.E.A Institute (Council for Agricultural Research and Economics) of Sanremo, Italy where the studied species were cultivated in greenhouses.

**Figure 2:**
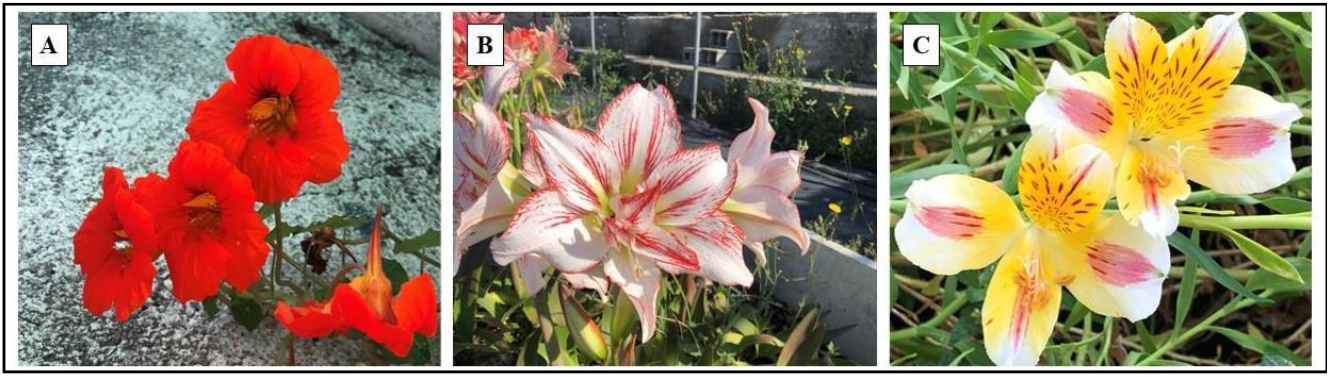
Images of the flowers of all species studied. (A) *Tropaeolum majus* L. (Fam.: Tropaeolaceae); (B) *Hippeastrum vittatum* Herb. (Fam.: Amaryllidaceae); (C) *Alstroemeria aurea* Graham (Fam.: Alstroemeriaceae).

### 2.3. Pollen sampling

Three pollen collection approaches were adopted (Table 1; Fig. 3): i) anthers were collected by carefully removing them from the flowers using forceps; ii) pollen grains obtained by dehisced anthers through multiple steps of sieving starting with a mesh of 100 μm size to a final mesh of 50 μm in order to isolate pollen grains from other floral parts and used as the control group (hereafter: mesh); (iii) E-PoSa developed in this study. For each sampling approach, we collected pollen from 20 flowers per species to gain enough material for all the subsequent analyses. All the collected samples were dried in an oven at 30°C for 12 hours. The three methods were compared for their recovery efficiency and the differences occurring in the macronutrient profiling.

**Table 1:**
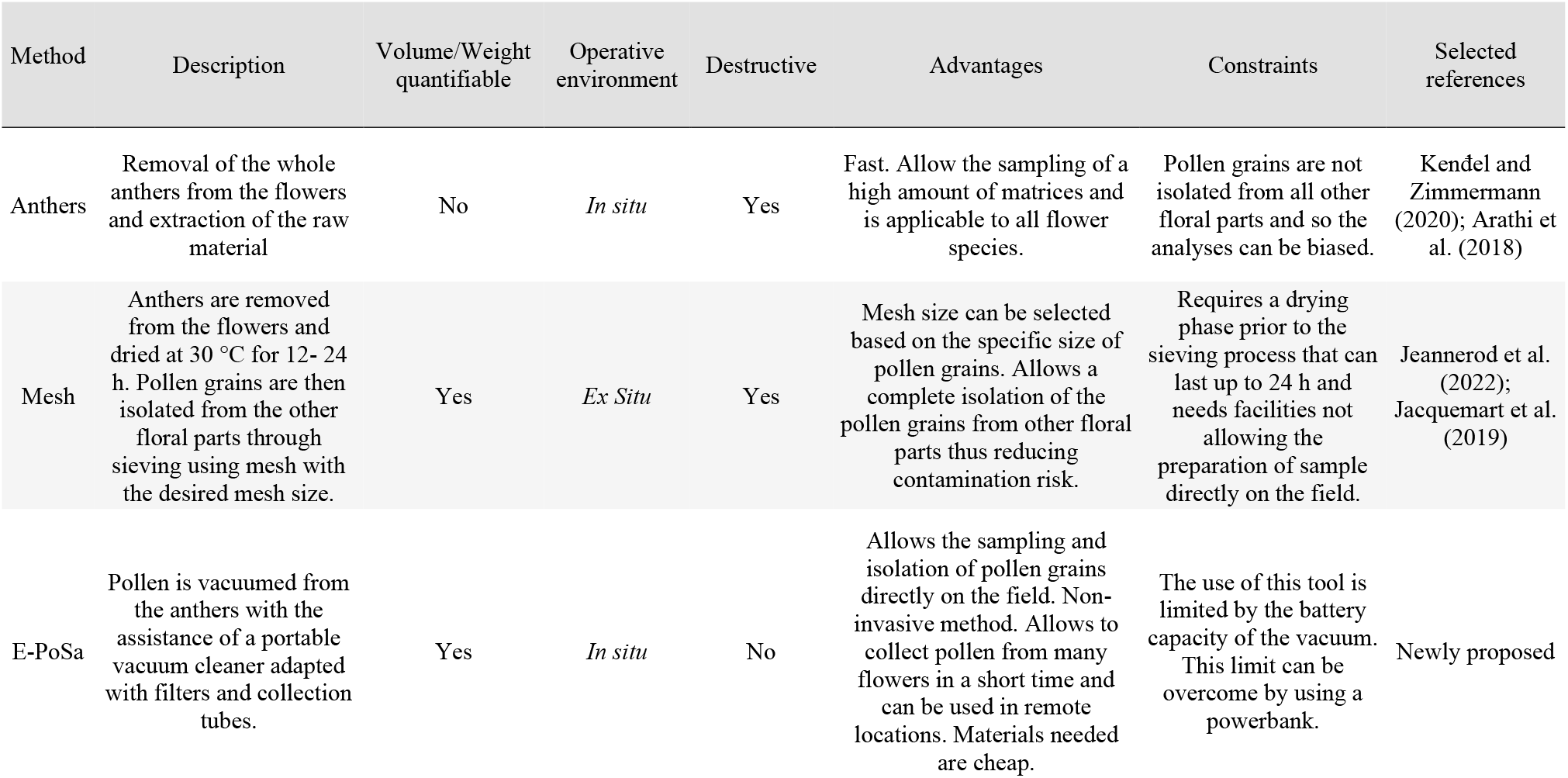
The table reports a brief description and the characteristics of the three methods used in this study for pollen sampling.

**Figure 3:**
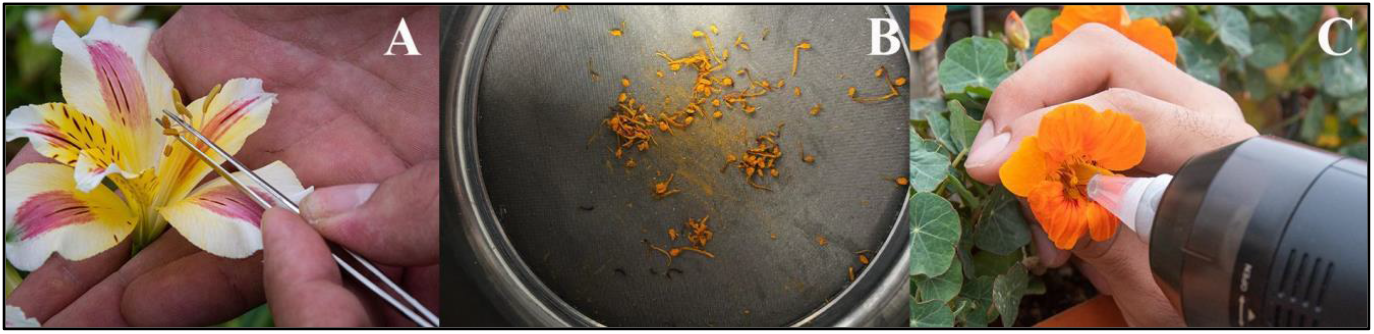
Photos of the different sampling techniques utilized for pollen collection: A. Removal of the anthers from *A. aurea* flower; B. Separation of pollen grains from the anthers of *H. vittatum* using a mesh; C. Pollen sampling with the E-PoSa device from *T. majus*.

Details on the nutritional analysis are reported in Appendix A1.

### 2.4. Statistical analysis

The differences in the recovery of pollen (expressed as mg per flower) were evaluated by a Linear Mixed Effects Model. The collection method was the fixed effect while the sampled species was accounted as a random factor. The software used was R (1.4.1106) and the package exploited was “nlme” (Pinheiro & Bates, 2023). For details on the statistical analysis on pollen nutritional composition see Appendix A1.

### 2.4. Validation in operative environment

The E-PoSa was employed also in the operative environment to collect pollen from a diverse array of wildflowers in the framework of a project aiming at the characterization of the pollinators nutritional landscape of northern Italian meadows. The sampling protocol adopted was the same as in the greenhouse validation experiment. The number of flowers/inflorescences sampled per species was recorded in the field and related to the weight of the pollen retrieved to estimate both the pollen recovery per floral unit and the minimum number of floral units to be sampled to perform nutritional analysis.

## 3. Results

Compared to other sampling approaches, the adoption of the E-PoSa represents a cost-effective and easy-to-use tool for the sampling of pollen from a wide range of flower species with different flower morphology and pollen yields. As anthers sampling does not allow measuring the weight of the pollen retrieved directly, it was not possible to compare the mass recovered by this technique with the mesh and the E-PoSa. Pollen collection through the E-PoSa device resulted in a significantly higher amount of pollen retrieved (0.68 ± 0.18 mg/flower) compared to the mesh method (0.27 ± 0.11 mg/flower) as shown in Fig. 4. The present method let us perform a field sampling on at least 22 wildflower species belonging to 8 families showing a broad range of flower morphology, that were successfully collected for the characterization of pollen nutrients (see Table 2 for details).

**Table 2:**
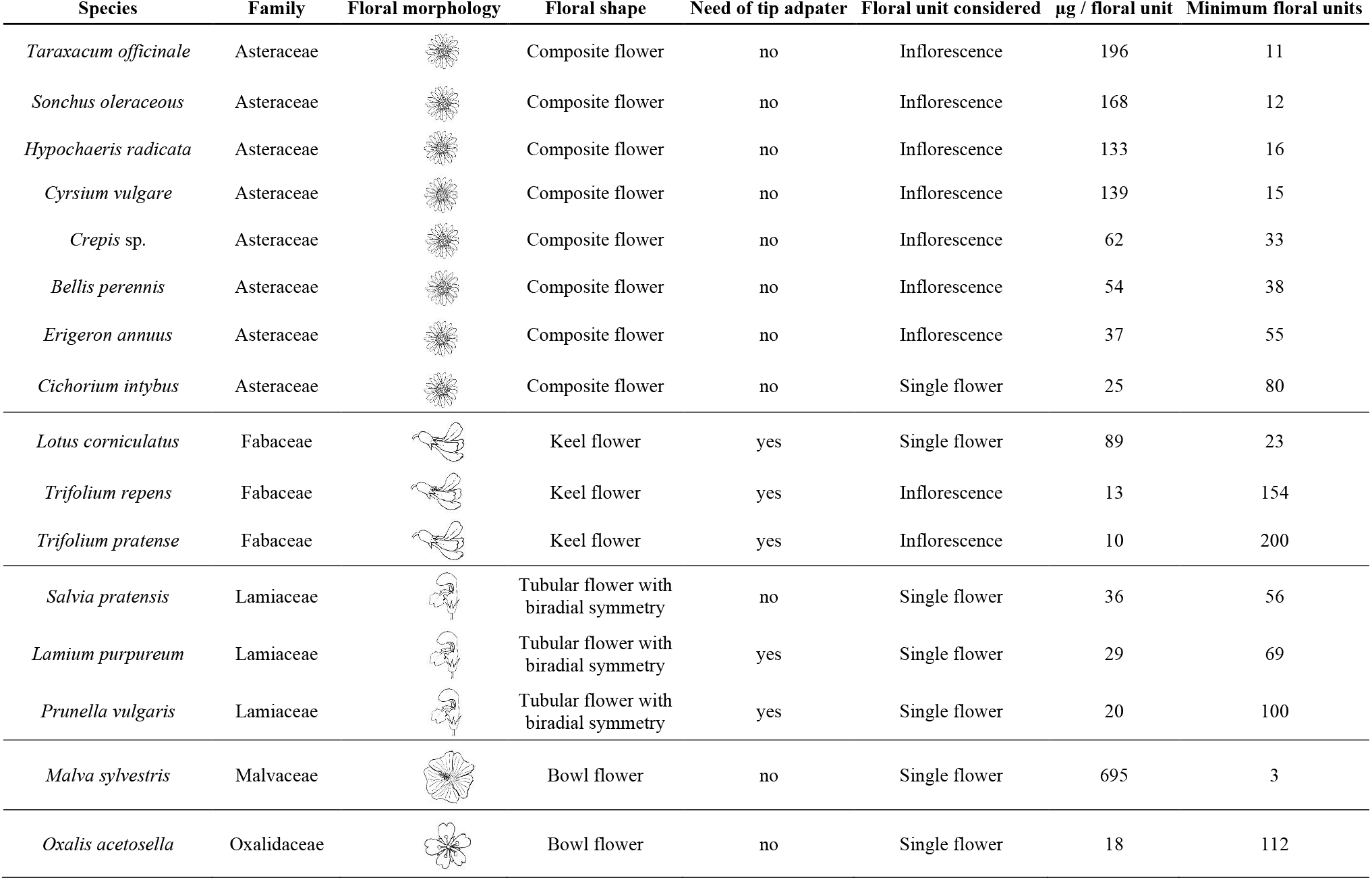
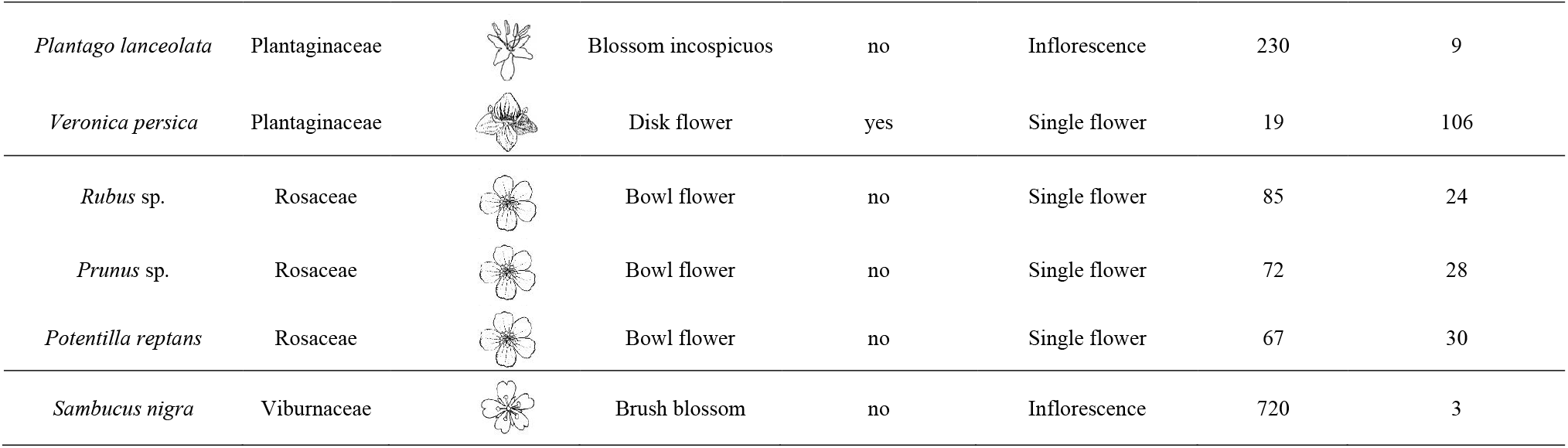
The table provides information on pollen recovery from different wildflower species. It includes family, floral morphology, and the need for E-PoSa adaptation with the tip. The “minimum floral units” column indicates the required number of floral units for obtaining 2 mg of pollen, the minimum amount for macronutritional profiling (proteins, lipids, and carbohydrates) using the protocol described in Appendix A1.

| Species | Family | Floral morphology | Floral shape | Need of tip adapter | Floral unit considered | µg / floral unit | Minimum floral units |
| --- | --- | --- | --- | --- | --- | --- | --- |
| *Taraxacum officinale* | Asteraceae |  | Composite flower | no | Inflorescence | 196 | 11 |
| *Sonchus oleraceus* | Asteraceae |  | Composite flower | no | Inflorescence | 168 | 12 |
| *Hypochaeris radicata* | Asteraceae |  | Composite flower | no | Inflorescence | 133 | 16 |
| *Cirsium vulgare* | Asteraceae |  | Composite flower | no | Inflorescence | 139 | 15 |
| *Crepis* sp. | Asteraceae |  | Composite flower | no | Inflorescence | 62 | 33 |
| *Bellis perennis* | Asteraceae |  | Composite flower | no | Inflorescence | 54 | 38 |
| *Erigeron annuus* | Asteraceae |  | Composite flower | no | Inflorescence | 37 | 55 |
| *Cichorium intybus* | Asteraceae |  | Composite flower | no | Single flower | 25 | 80 |
| *Lotus corniculatus* | Fabaceae |  | Keel flower | yes | Single flower | 89 | 23 |
| *Trifolium repens* | Fabaceae |  | Keel flower | yes | Inflorescence | 13 | 154 |
| *Trifolium pratense* | Fabaceae |  | Keel flower | yes | Inflorescence | 10 | 200 |
| *Salvia pratensis* | Lamiaceae |  | Tubular flower with<br>biradial symmetry | no | Single flower | 36 | 56 |
| *Lamium purpureum* | Lamiaceae |  | Tubular flower with<br>biradial symmetry | yes | Single flower | 29 | 69 |
| *Prunella vulgaris* | Lamiaceae |  | Tubular flower with<br>biradial symmetry | yes | Single flower | 20 | 100 |
| *Malva sylvestris* | Malvaceae |  | Bowl flower | no | Single flower | 695 | 3 |
| *Oxalis acetosella* | Oxalidaceae |  | Bowl flower | no | Single flower | 18 | 112 |
| *Plantago lanceolata* | Plantaginaceae |  | Blossom incospicuos | no | Inflorescence | 230 | 9 |
| *Veronica persica* | Plantaginaceae |  | Disk flower | yes | Single flower | 19 | 106 |
| *Rubus* sp. | Rosaceae |  | Bowl flower | no | Single flower | 85 | 24 |
| *Prunus* sp. | Rosaceae |  | Bowl flower | no | Single flower | 72 | 28 |
| *Potentilla reptans* | Rosaceae |  | Bowl flower | no | Single flower | 67 | 30 |
| *Sambucus nigra* | Viburnaceae |  | Brush blossom | no | Inflorescence | 720 | 3 |

**Figure 4:**
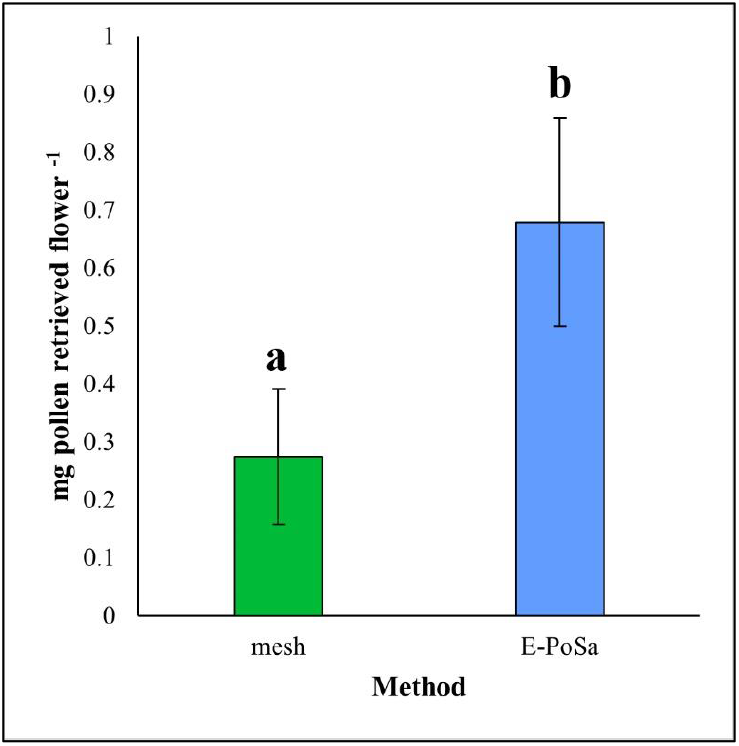
Barplot reports the mg of pollen retrieved per flower depending on the sampling method. The results are shown as the mean values obtained for the three species sampled. Values are reported as the mean ± SEM (standard error of the mean). Significant differences (*p* < 0.05) estimated through the Linear Mixed Model are reported by different letters.

### 3.1. Pollen macronutrient composition

Estimation of the macronutrient composition of pollen is shown in Fig. 5. The total protein content was significantly lower in dry anthers compared to E-PoSa and mesh (Table S2). No significant differences emerged in the comparison between the total protein content in pollen collected with E-PoSa and the mesh. The lipid content was significantly lower in anthers than in E-PoSa while in mesh samples it was comparable both to E-PoSa and the anthers (Table S2). Concerning the sugar composition of pollen, glucose was significantly higher in the anthers compared both to E-PoSa and the mesh control group (Table S2); similarly, fructose content was higher in anthers compared to the mesh while E-PoSa did not significantly differ with anthers and mesh (Table S2). Finally, the sucrose content was not significantly different among the three sampling methods (Table S2).

**Figure 5:**
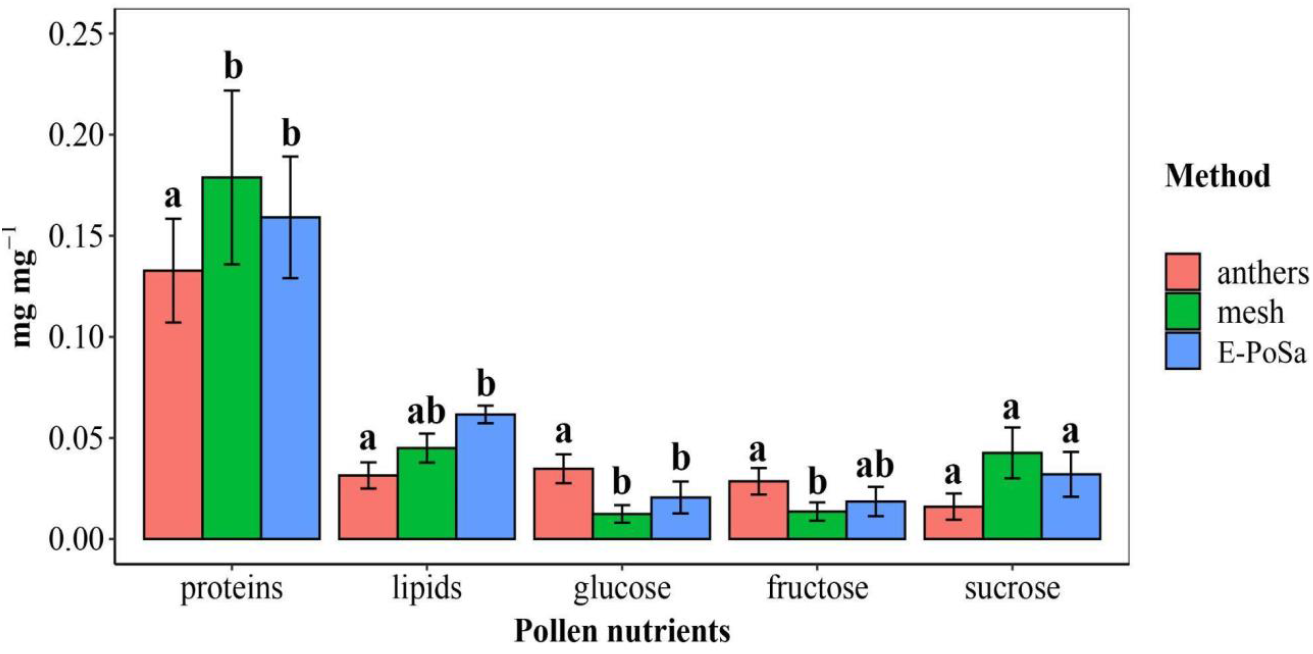
Quantified percentages of the macronutrient composition of the samples in the 3 species studied, which are implemented into the regression model as a random factor. The results are shown as the mean values obtained for the three species sampled. Values are reported as the mean ± SEM (standard error of the mean). For each of the detected analytes, significant differences (*p* < 0.05) estimated through the post hoc Tukey test are declared within the same nutritional category and are reported by different letters.

## 5. Discussion

The recent advances in pollination nutritional ecology require the definition of standardized sampling methods for pollen (Jeannerod et al., 2022). The analysis of pollen from wild plants needs to be fine-tuned for an accurate definition of its role in the nutritional balance of pollinators’ diets (Lau et al., 2022). It is well known that the recovery of pollen grains from wild plants is a difficult task to perform since the anthers of enthomogamous plant species usually produce low amounts of pollen (e.g., Jeannerod et al., 2022; Palmer-Young et al., 2019). Obtaining a significant amount of pollen without contamination originating from other floral parts can be very time-consuming, reducing the efficiency and the feasibility of nutritional ecology studies. The results of the nutritional analyses of pollen performed on the three target species show that this strategy avoids sample contaminations since the pollen collected by the E-PoSa did not show any significant differences in terms of macronutrient composition compared to the pollen sampled by sieving. In this framework, we suggest the adoption of the E-PoSa for the non-invasive collection of a satisfactory amount of pollen in a relatively short time. The adoption of the E-PoSa does not require any expertise or training for its usage due to the ease of assembly and the getting of the equipment. Furthermore, it lets the sampling of pollen free of contaminants possibly impairing further nutritional analyses. This feature makes it possible to conceive its exploitation in multiple contexts such as in the framework of citizen science activities, plant breeding programmes, and/or in contexts requiring the need to optimize the logistics and human resources available. The present research is to be intended as the precursor of future nutritional ecology studies dealing with the characterization of pollen chemical composition of plant communities consisting of a broad range of floral morphologies. The effectiveness of the recovery allowed the sampling to be performed on a wide range of species considering the availability of floral resources usually occurring in meadows (see Table 2).

## Conclusions

The field of nutritional ecology in the context of plant-pollinator interactions is growing in importance to address the issues of pollinator safeguarding and conservation. The novel E-PoSa can provide a cheap and easy-to-assemble tool, encouraging its future use in the field of pollen nutrition.

## Award Number

Project code CN_00000033, Concession Decree No. 1034 of 17 June 2022 adopted by the Italian Ministry of University and Research, CUP, H43C22000530001 Project title “National Biodiversity Future Center - NBFC”.

L.T. is supported by the PhD Program PON “Ricerca e Innovazione” 2014-2020, DM n. 1061 (10/08/2021) and n. 1233 (30/07/2020).

The authors are grateful to Barbara Ruffoni, Paolo Mussano and the team of the CREA of Sanremo for their support. A special thank goes to Fausto Ramazzotti for his help during the sampling activity.

## AUTHOR CONTRIBUTIONS

EP, LC, LT and AG conceived the ideas and designed the methodology; EP, LG and AC collected the data; EP and LG analyzed the data; EP, LG, PB and AG led the writing of the manuscript. All authors critically contributed to the draft and gave their final approval for publication.

## DATA AVAILABILITY

All data can be accessed at the following FigShare link: https://figshare.com/s/db777fb95f1056c20843

