## Supplementary information for "E-PoSa: a novel and effective tool for sampling pollen directly from flowers"

**Table S1**

| **Item** | **Provider** | **Cost (€)** | **Note** |
| --- | --- | --- | --- |
| Mini-USB Vacuum Cleaner | Honk (Amazon) | 22.99 |  |
| Inox mesh 30 x 100 cm. 0,075 mm aperture | TIMESETL (Amazon) | 16.99 |  |
| Power Bank 26800mAh, Portable charger USB C Powerbank Battery | Charmast (Amazon) | 34.99 | Optional |

**Table S1:** Price of the necessary hardware for E-PoSa assembly.

**Table S2**

| **Macronutrient** | **Method** | **Mean content (% *w/w*)** | **SEM** |
| --- | --- | --- | --- |
| Proteins | Anthers | 13.27 | 3.8 |
|  | Mesh | 17.77 | 3.82 |
|  | E-PoSa | 15.92 | 3.63 |
| Lipids | Anthers | 3.15 | 0.94 |
|  | Mesh | 4.87 | 0.8 |
|  | E-PoSa | 6.63 | 0.64 |
| Glucose | Anthers | 3.48 | 1.16 |
|  | Mesh | 1.24 | 0.43 |
|  | E-PoSa | 2.1 | 0.79 |
| Fructose | Anthers | 2.82 | 0.65 |
|  | Mesh | 1.36 | 0.72 |
|  | E-PoSa | 1.86 | 0.45 |
| Sucrose | Anthers | 1.6 | 0.65 |
|  | Mesh | 4.26 | 0.93 |
|  | E-PoSa | 3.2 | 1.11 |

**Table S2:** Details on the nutritional composition of pollen collected with the three different methods. The values are reported as the mean values obtained for the three species sampled ± SEM (standard error of the mean).

**Video S1**

https://clipchamp.com/watch/0uBKbA8kYUei
